# Whole-genome reference panel of 1,781 Northeast Asians improves imputation accuracy of rare and low-frequency variants

**DOI:** 10.1101/600353

**Authors:** Seong-Keun Yoo, Chang-Uk Kim, Hie Lim Kim, Sungjae Kim, Jong-Yeon Shin, Namcheol Kim, Joshua SungWoo Yang, Kwok-Wai Lo, Belong Cho, Fumihiko Matsuda, Stephan C. Schuster, Changhoon Kim, Jong-Il Kim, Jeong-Sun Seo

## Abstract

Genotype imputation using the reference panel is a cost-effective strategy to fill millions of missing genotypes for the purpose of various genetic analyses. Here, we present the Northeast Asian Reference Database (NARD), including whole-genome sequencing data of 1,781 individuals from Korea, Mongolia, Japan, China, and Hong Kong. NARD provides the genetic diversities of Korean (n=850) and Mongolian (n=386) ancestries that were not present in the 1000 Genomes Project Phase 3 (1KGP3). We combined and re-phased the genotypes from NARD and 1KGP3 to construct a union set of haplotypes. This approach established a robust imputation reference panel for the Northeast Asian populations, which yields the greatest imputation accuracy of rare and low-frequency variants compared with the existing panels. Also, we illustrate that NARD can potentially improve disease variant discovery by reducing pathogenic candidates. Overall, this study provides a decent reference panel for the genetic studies in Northeast Asia.

---

During the past decade, whole-genome sequencing (WGS) of the reference populations has enabled the extensive human genetic researches to be carried out^1,2^. It has played an imperative role in the genetic researches, especially for genotype imputation in the genome-wide association study (GWAS). With the recent expanding number of such studies, several research groups have generated the extensive WGS data to build reference panels^1–10^. The most commonly used imputation panels are constructed by the 1000 Genomes Project Phase 3 (1KGP3) and Haplotype Reference Consortium (HRC), which are publicly available for researchers. As genotype imputation increases the power of GWAS in a cost-efficient way, the confidence of imputed genotypes is important. To improve the quality of imputation in the genetic studies, large-scale population-specific panels with deep-coverage WGS need to be used^4,9^.

Despite Northeast Asians account for 21.51% of worldwide population (see URLs), the majority of genetic studies and the reference panels have European bias^11^. There are only a few population-scale WGS studies covering Northeast Asians from China, Japan, and Mongolia^6,8,12,13^, and these studies have the several issues for the improved reference panel in Northeast Asia such as public unavailability, inadequate sequencing coverage, and small sample size. Furthermore, although Koreans (KOR) are one of the major population groups in Northeast Asia, previous datasets for KOR^14–16^ does not have enough number of WGS samples to accurately impute the genome-wide variants of KOR population. Therefore, constructing a large-scale whole-genome reference panel for the diverse population groups in Northeast Asia with deep sequencing coverage is still necessary to allow dense and accurate genotype imputation for the genetic researches in these populations.

In this study, we constructed the Northeast Asian Reference Database (NARD), consisting of 1,781 individuals from Korea, Japan, Mongolia, China, and Hong Kong. The goal of this study is to establish a high-quality population-specific reference panel for the genetic studies and precision medicine in Northeast Asia.

## Results

### Variants statistics

WGS was performed on 1,781 Northeast Asians including KOR (n = 850), Japanese (JPN, n = 396), Mongolian (MNG, n = 386), Han Chinese (CHN, n = 91), and Hong Kongese (HKG, n = 58) with deep (n = 834, ≥ 20X) or intermediate (n = 947, 10X~20X) sequencing coverages (Supplementary Figure 1 and Supplementary Table 1).

In NARD, a total of 40.6 million single nucleotide polymorphisms (SNPs) and 3.8 million short insertions/deletions (indels) were discovered, and 77.1% were singletons or rare variants (minor allele frequency [MAF] < 0.5%; Table 1). On average, 3.3 million SNPs and 0.3 million indels were found per individual. We identified 15.4 million novel SNPs (37.8% of the total) in NARD (Supplementary Fig. 2a). Among them, 45.0% were specific to KOR, likely due to their large sample size, and 12.6% were found across whole populations in NARD (Supplementary Fig. 2b). The majority of novel SNPs were singleton or rare, and located in the non-coding regions (Supplementary Fig. 2c). In order to measure the integrity of our WGS variant call pipeline, we calculated the genotype concordance between WGS and Illumina Omni 2.5M array of 86 CHN samples in NARD. The genotype concordance between two datasets was 99.6% (Supplementary Table 2), reflecting high integrity of our pipeline and high confidences of our variants.

**Table 1.** Total number of variants in 1,781 East Asians by MAF and functional category.

| Variant type | Frequency<sup>a</sup> | Number of variants | Functional variation |  |  |  |  |  |  |  |  |
| --- | --- | --- | --- | --- | --- | --- | --- | --- | --- | --- | --- |
|  |  |  | Protein coding region |  |  |  | Non-coding region |  |  |  |  |
|  |  |  | Silent / Nonframeshift | Missense / Frameshift | Stoploss / Stopgain | Unknown | Intronic | Intergenic | Splicing | UTR | ncRNA |
| SNP | Singleton | 17,732,020 | 86,641 | 146,235 | 3,714 | 2,685 | 6,824,083 | 9,314,177 | 2,109 | 246,964 | 1,105,412 |
|  | Rare | 13,776,881 | 54,844 | 88,130 | 1,670 | 1,923 | 5,293,510 | 7,322,759 | 1,367 | 165,066 | 847,612 |
|  | Low | 3,428,747 | 12,777 | 15,692 | 230 | 428 | 1,300,329 | 1,848,843 | 244 | 38,718 | 211,486 |
|  | Common | 5,711,192 | 17,853 | 15,957 | 151 | 729 | 2,046,271 | 3,216,818 | 159 | 53,127 | 360,127 |
|  | Total | 40,648,840 | 172,115 | 266,014 | 5,765 | 5,765 | 15,464,193 | 21,702,597 | 3,879 | 503,875 | 2,524,637 |
| Indel | Singleton | 1,398,360 | 3,189 | 5,067 | 157 | 129 | 557,813 | 714,077 | 522 | 27,709 | 89,697 |
|  | Rare | 1,387,257 | 2,752 | 2,898 | 130 | 127 | 546,652 | 724,226 | 217 | 22,143 | 88,112 |
|  | Low | 451,268 | 640 | 825 | 34 | 38 | 173,863 | 240,553 | 61 | 6,434 | 28,820 |
|  | Common | 567,341 | 419 | 366 | 18 | 89 | 206,814 | 315,440 | 145 | 7,142 | 36,908 |
|  | Total | 3,804,226 | 7,000 | 9,156 | 339 | 383 | 1,485,142 | 1,994,296 | 945 | 63,428 | 243,537 |

<sup>a</sup> Rare; MAF < 0.5%, low; 0.5% ≤ MAF < 5% common; MAF ≥ 5%.

### Ancestry composition of NARD

We examined the ancestry composition of the individuals in NARD by conducting population structure analyses using NARD combined with 1KGP3 data, to illustrate how NARD covers the genetic diversities that were not present in other panels. From the result of principal component analysis (PCA) of global human populations, the individuals from NARD were closely related to East Asians from 1KGP3 as expected. MNG were separately clustered and positioned between East Asian and non-African populations as previously reported^8^ (Fig. 1a). When we applied PCA to only East Asians, a clear population differentiation pattern was observed in NARD (Fig. 1b); MNG was most distinct from other populations based on PC1, and PC2 separated KOR, JPN, and mainland East Asians (CDX, CHB, CHS, HKG, and KHV). Interestingly, KOR and JPN were not overlapped each other except for some outliers. This underlies that their ancestral compositions are distinctive enough to form the separate clusters. Additionally, the unsupervised ADMIXTURE analysis^17^ supported the different ancestral components for each of KOR, MNG, JPN, and mainland East Asians (Fig. 1c). The results highlight that NARD represents the diverse genetic compositions within Northeast Asian populations, by including the two ancestries, KOR and MNG, which have been underrepresented in public reference panels such as 1KGP3.

**Fig. 1.**
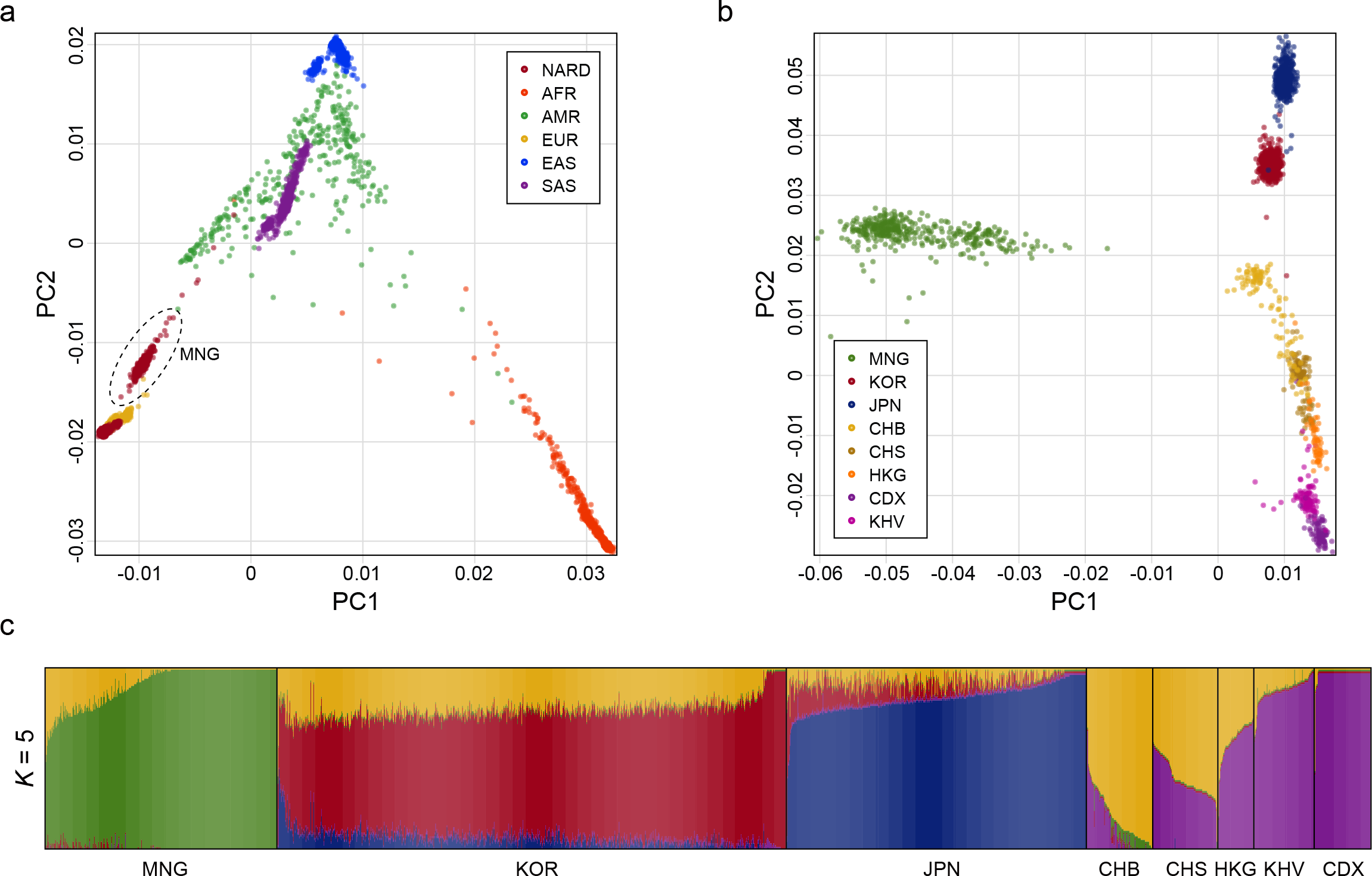
Ancestry composition of 1,781 individuals in NARD. **a,** Principal component analysis (PCA) of NARD and the global populations from 1KGP3. **b,** PCA of NARD and East Asians from 1KGP3. JPT from 1KGP3 were combined into JPN. CHN of NARD were categorized into CHB and CHS. **c,** Population substructure of East Asians with five ancestral components inferred by ADMIXTURE algorithm.

### Evaluation of NARD imputation panel

To illustrate the robustness of NARD as an imputation reference panel, we built a pseudo-GWAS dataset using an independent cohort of 97 unrelated KOR individuals^14,18,19^. It was generated from WGS data by masking the genotypes that were not included in the sites of Illumina Omni 2.5M array. These sites were also used for the imputation accuracy measurement in the previous studies of Northeast Asians^6,8^. Then, the imputation was conducted by Minimac3^20^ on pre-phased SNPs using five types of reference panel: 1) NARD (n = 1,781), 2) 1KGP3 (n = 2,504), 3) HRC r1.1 (n = 32,470), 4) NARD + 1KGP3 (n = 4,202), and 5) NARD + 1KGP3 (re-phased, n = 4,202). To measure the imputation accuracy, we calculated the squared Pearson correlation coefficients (*R*^2^) between the true genotypes and the imputation dosages as a function of MAF for 850 KOR from NARD. The imputation performance of NARD exceeded 1KGP3 for every MAF bins (Fig. 2a). Notably, HRC panel, with the largest sample size comprising 1KGP3, showed poor imputation result compared with other panels.

**Fig. 2.**
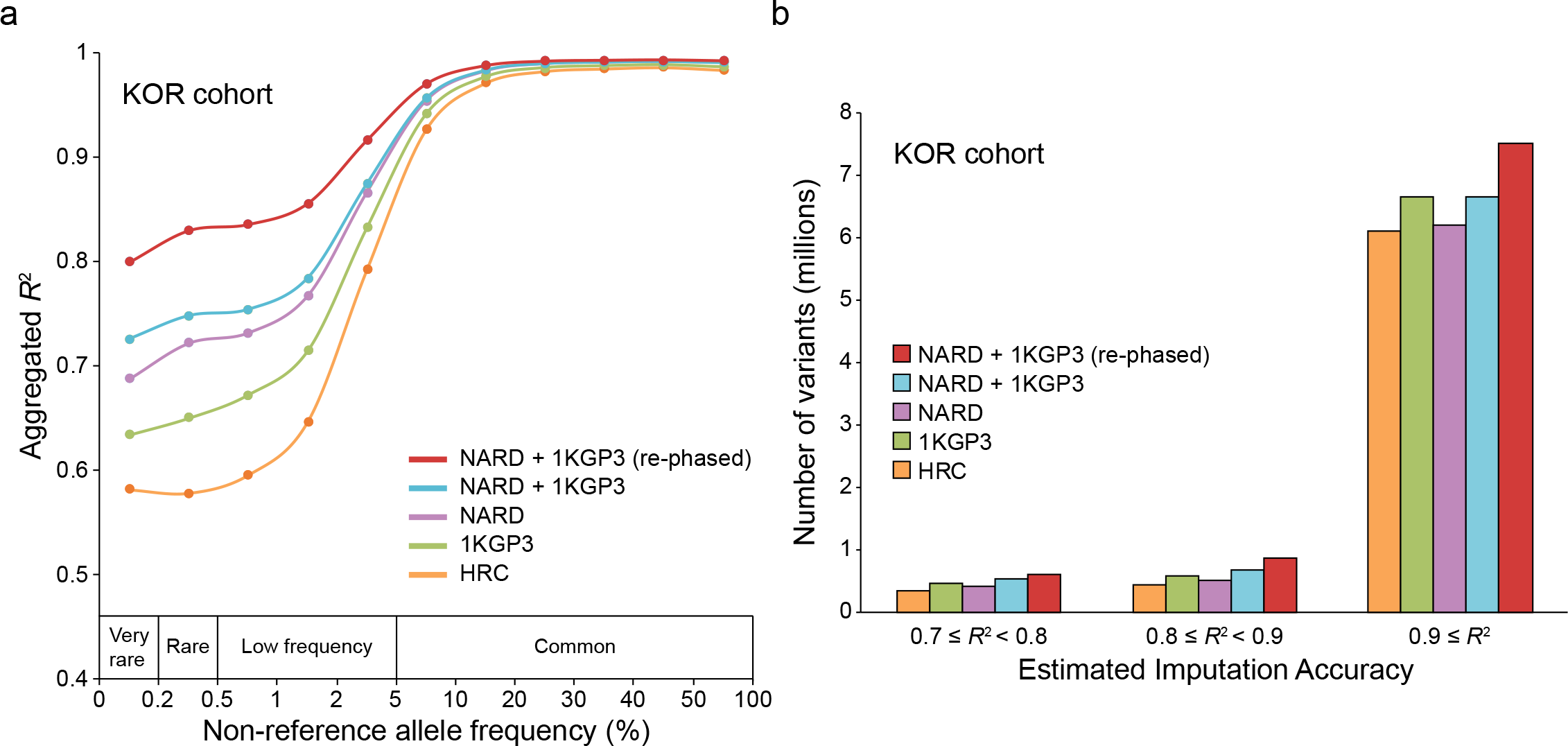
Imputation performance evaluation. **a,** Imputation accuracy assessment using five different reference panels. The pseudo-GWAS panel of 97 KOR was used for imputation. The *x*-axis represents MAF of 850 KOR individuals from NARD. The *y*-axis represents the aggregated *R*^2^ values of SNPs, which were calculated by the true genotypes and the imputed dosages. Only SNPs that were imputed across all panels were used for the aggregation of *R*^2^ values. **b**, Number of imputed SNPs as a function of the estimated imputation accuracy and the types of imputation panel. This result was generated based on the *R*^2^ values that were estimated by Minimac3.

We then merged NARD and 1KGP3 panels, and performed re-phasing to further enhance the imputation performance based on the previous studies^2,5^. To combine NARD and 1KGP3 without missing genotypes, we used the identical approach as implemented in UK10K^5^ and IMPUTE2^21^ (see URLs); we reciprocally imputed two panels using Minimac3, therefore missing genotypes in NARD or 1KGP3 were statistically inferred. Consistent with the previous studies^4,6,8–10^, combining two panels showed more accurate imputation result compared with NARD or 1KGP3 alone. Furthermore, we confirmed a large improvement of the imputation accuracy, particularly for very-rare (MAF < 0.2%; *R*^2^ = 0.80), rare (0.2% ≤ MAF < 0.5%; *R*^2^ = 0.83), and low-frequency (0.5% ≤ MAF < 5%; *R*^2^ = 0.87) variants, when the haplotypes in the combined panel were re-phased by SHAPEIT3^22^. In addition to measuring accuracy, we assessed the number of the accurately imputed SNPs for each panel. For this analysis, we used the estimated *R*^2^ values in the info file measured by Minimac3, as it is the standard for the quality control procedure in GWAS^23,24^. We found that the re-phased panel produced the greatest number of high-confident SNPs (*R*^2^ ≥ 0.9) compared with other panels, especially with 1KGP3 (n = 7.5 million versus 6.7 million), in concordance with the imputation accuracy (Fig. 2b).

To investigate the underlying reason of the improved imputation performance by the re-phasing approach, we performed the identity-by-descent (IBD) analysis. It is known that phasing or genotype errors cause the gaps within the real IBD tracts, hence shorter segments tend to be detected in phased genotype data^25,26^. Based on this aspect, we expected that haplotype correction is occurred in NARD by re-phasing and it would extend the length of shared IBD segments among individuals. Therefore, we measured the shared large IBD segments (≥ 2 cM) between the two individuals using the original (phased without 1KGP3) and the re-phased haplotypes of NARD. As a result, we confirmed the significant increase of shared IBD lengths and numbers in the re-phased haplotypes, which implies that the haplotype refinement is applied to NARD by the long-range phasing (LRP)^27,28^ during the re-phasing process (Supplementary Fig. 3).

We further represented the power of the re-phased panel using an independent cohort of 106 unrelated Northeast Asian individuals (79 CHN and 27 JPN)^29,30^. For the imputation accuracy measurement, we used MAF bins defined by 2,093 Northeast Asians from NARD and 1KGP3 (CHB, CHS, and JPT). In agreement with the imputation result for the KOR cohort, the re-phased panel provided the most accurate genotype dosages on very-rare (*R*^2^ = 0.74), rare (*R*^2^ = 0.75), and low-frequency (*R*^2^ = 0.81) variants (Supplementary Fig. 4a). Moreover, the re-phased panel also generated the largest number of accurate imputed genotypes compared with other panels, especially with 1KGP3 (n = 7.3 million versus 7.0 million; Supplementary Fig. 4b).

### NARD imputation server

We have developed a web-based imputation server for the researchers to publicly use NARD + 1KGP3 (re-phased) panel (see URLs). The NARD imputation server provides the imputation process for a wide range of genotype data format including PLINK^31^ (ped files paired with map files or bed files paired with bim and fam files), 23andMe (Mountain View, CA), AncestryDNA (Lehi, UT) files, and the variant call format (VCF). The results are processed through the imputation pipeline consisting of four major steps; pre-processing, phasing, imputation, and post-processing (Supplementary Fig. 5). The pre-processing step checks the uploaded files in valid formats and converts them into VCF files for the next steps. If the input data is uploaded in a PLINK format, it will be converted based on human_g1k_v37 reference from 1KGP3 using GotCloud^32^, and 23andMe/AncestryDNA files will be converted by bcftools^33^. The pre-processed data is phased using Eagle or SHAPEIT2^34^ with or without a reference panel, respectively, as the pipeline of Michigan Imputation Server (see URLs). Then, imputation is implemented with Minimac3. In the post-processing step, the output is assessed and provided as the gzip-compressed VCF files.

### NARD for variant interpretation

We also evaluated the advantage of NARD as a population-specific reference panel for the clinical variant interpretation. Common variants were excluded for this analysis because it is the first step to identify rare disease-causing genes^35^. To examine the potential advantage of NARD, the frequencies of SNPs between the Genome Aggregation Database (gnomAD 2.1.1 release; see URLs) and NARD were compared. We redefined the frequency of 1.8 million genome-wide SNPs that are rare in worldwide populations from gnomAD (gnomAD-ALL) to low-frequency or common (MAF ≥ 5%). Moreover, 0.9 million rare genome-wide SNPs in East Asian from gnomAD (gnomAD-EAS) were low-frequency or common variants in NARD (Fig. 3a).

**Fig. 3.**
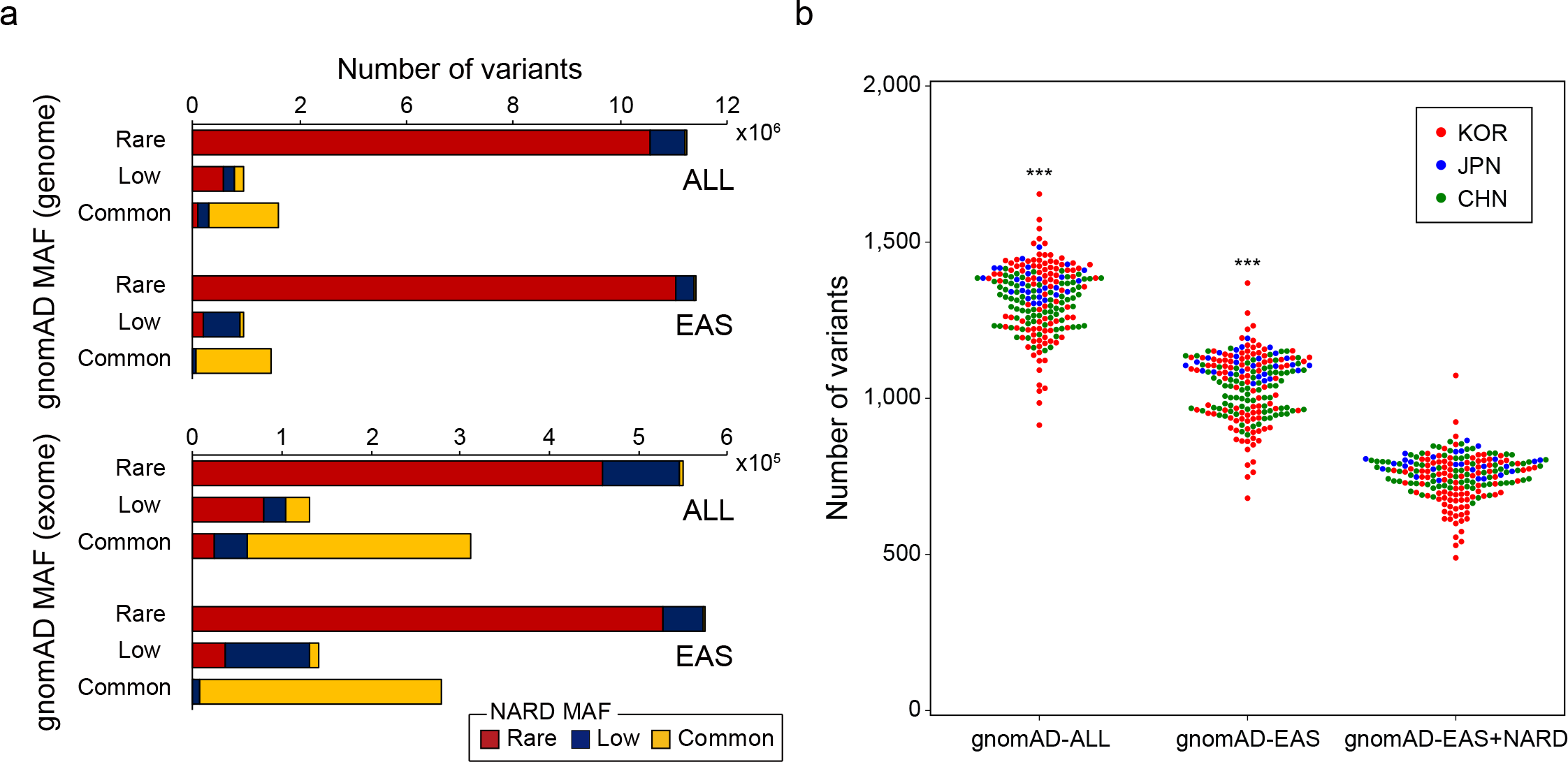
Variant interpretation using NARD. **a,** Allele frequency differences of SNPs shared between NARD and gnomAD. The *x*-axis denotes the MAF of SNPs in the global populations (ALL) or East Asians (EAS) from gnomAD database. Color represents the MAF of SNPs in 1,781 Northeast Asians from NARD. **b,** Number of uncommon (MAF < 5%) protein-altering variants (missense, nonsense, frameshift, and splicing variants) after filtration according to each database: gnomAD-ALL, gnomAD-EAS, and gnomAD-EAS + NARD. Variant catalogue from gnomAD exome was applied. Common variants were filtered using the gnomAD with or without NARD. \*\*\**P* < 0.0001 by two-tailed Mann-Whitney U-test (compared with gnomAD EAS + NARD).

Then, we simulated rare disease variant discovery using 203 samples that were included in two pseudo-GWAS panels for the imputation analysis. We applied the variant filtering criteria (MAF < 5%) from the guideline of American College of Medical Genetics for the interpretation of sequence variants^36^. Notably, the number of protein-altering variants (missense, nonsense, frameshift, and splicing variants) was significantly reduced when the exome catalogue of gnomAD-EAS and NARD were jointly applied for variant filtration (Fig. 3b). This result represents that NARD could also contribute to inference of the pathogenic variant classification as well as genotype imputation for the Northeast Asians.

## Discussion

Due to the cost-reduction and the technological advancements in WGS, several groups have been focused on building the population-specific reference panels, especially for underrepresented populations in the conventional panels such as 1KGP3^3,4,7–10^. However, the Northeast Asian-specific panel with deep sequencing coverage and large sample size has been barely constructed. In this study, we integrated whole-genome sequence variants of 1,781 Northeast Asian individuals to construct a reference panel, NARD, to resolve the uncertainty in genotype imputation using the pre-existing panels and to facilitate more comprehensive genetic researches in Northeast Asia.

Genotype-imputation accuracy is known to be affected by several factors and one of the major determinants is the size of reference panel^5,34^. Hence, most genetic studies for the Northeast Asians^37–40^ were relied on the panels with large sample size, although the ancestries between the study population and the reference panel are not matched. However, these panels showed lower power in genotype imputation, compared to the well-matched population-specific panels with smaller sample size^4,6–10,41^. Recently, HRC panel was constructed using the genotypes of more than 30,000 individuals (mostly Europeans), but previous study demonstrated the poor performance of this panel for the Northeast Asians, even worse than 1KGP3 panel^42^, and our analysis showed the same result. This might be due to the skewed proportion of European ancestry in HRC panel as unhelpful haplotypes for the Northeast Asians^43^, and emphasizing the importance of the population-specific reference panel for the accurate genotype imputation and the subsequent genetic analysis. Considering the importance of the population-specific reference panel, we generated large-scale WGS dataset of KOR and MNG that were not included in 1KGP3. From the results of population structure analysis, KOR and MNG were genetically differentiated from other East Asians. Therefore, the major ancestries in Northeast Asia are finally covered as population-scale by our dataset. In addition to two populations, other Northeast Asians including JPN, CHN, and HKG were also sequenced to increase the imputation power by sample size effect and to build NARD as a reference panel for the Northeast Asians.

As previous studies yield high imputation accuracy from their population-specific panels by combining dataset of 1KGP3^4,6,8–10^, we also confirmed the improvement of the imputation performance by combining NARD and 1KGP3 using a fast and simple approach as described in UK10K and IMPUTE2. However, there could be an issue regarding the uncertainty of the imputed genotypes, since the missing genotypes in each panel were statistically estimated. Therefore, referring to HRC study, calculating genotype likelihood of each variants using the individual BAM files in the panel would resolve this issue, if the sequencing coverages are sufficient. In addition to merging NARD and 1KGP3, we further enhanced the power of the combined panel by applying the re-phasing strategy. It is an advanced process that has not been applied in most previous studies^4,6,8–10^, but HRC study has shown further improvement of the imputation accuracy with this approach. Based on this strategy, NARD + 1KGP3 (re-phased) panel produced more accurate genotype dosages, especially for uncommon variants (MAF < 5%), than NARD + 1KGP3 panel, might be due to haplotype correction by LRP with the assistance of the haplotypes in 1KGP3 panel.

In summary, we generated a large-scale reference panel for the Northeast Asians, which will be a highly valuable resource to resolve the persistent deficiency of Asian genome data. We believe that our efforts will facilitate more extensive genetic researches and remarkably contribute to precision medicine in Northeast Asians.

## Methods

### Ethics statement

This study was approved by the institutional review board of Seoul National University Hospital, in accordance with the Declaration of Helsinki (approved ID: C-1705-048-852). Written informed consents were obtained from all study subjects.

### Whole-genome sequencing

Of 1,781 Northeast Asians, 1,690 individuals of KOR, JPN, MNG, and HKG, we performed WGS using Hiseq X instrument (Illumina, San Diego, CA) based on the manufacturer’s instruction. The others were from publicly available 91 CHN samples^43^, which were sequenced by Illumina Hiseq 2000 instrument (Illumina, San Diego, CA). This cohort consists of YH cell line, HapMap, and 1KGP3 samples with high sequencing depth (on average, 70X)^1,44,45^.

### Variant discovery and refinement

Read alignment to the human reference genome (hg19), duplicate read removal, and joint calling of SNPs/indels were performed using Dynamic Read Analysis for GENomics (DRAGEN) platform (version 01.003.024.02.00.01.23004; see URLs). For indels, we discarded variants greater than 49 base pairs. Variant quality score recalibration (VQSR) was applied to raw variants based on the GATK’s best practice^46^ (see URLs). SNPs and indels below 99% of truth sensitivity level from VQSR were initially filtered. Moreover, recalibrated variants were further filtered based on the following criteria: i) located in the low complexity regions (LCRs), which were defined by 1KGP3, ii) genotype quality < 20, and iii) read depth < 5. After several filtration processes, SNPs and indels were phased using SHAPEIT3 (version r884.1) using the default parameters without the reference panel.

### Variant discovery evaluation

We used the 86 CHN samples that simultaneously exist in NARD and Illumina Omni 2.5M array data downloaded from 1KGP3 site (see URLs). We excluded mitochondrial DNA and pseudoautosomal regions (PARs) to remain 1,664,330 SNPs overlapping between NARD and Omni chip for calculating the genotype concordance between two datasets. The concordance is the cumulative sum of the matching alleles divided by the total number of loci multiplied by two, which is the maximum matching opportunity in diploid. The male sex chromosomes are considered as diploids for this calculation.

### Variant annotation

All the SNPs and indels in this study were annotated by ANNOVAR^47^ based on RefSeq gene definition^48^. For novel variant classification, Kaviar^49^, gnomAD, and The Single Nucleotide Polymorphism Database build 150 (dbSNP)^50^ were annotated. To perform the simulation of rare disease variant discovery, we annotated MAF from gnomAD-All, gnomAD-EAS, and NARD on WGS data of 203 Northeast Asian individuals for imputation accuracy evaluation. Researchers could download MAFs of variants in NARD as ANNOVAR format (see URLs).

### PCA and ADMIXTURE analysis

We converted VCF files of bi-allelic autosomal SNPs from NARD and 1KGP3 into PLINK format using GotCloud (version 1.75.5). Then, we merged two panels by PLINK (version 1.9), and extracted SNPs with genotype rate equals to 100% and MAF ≥ 1% to remove the batch effect between NARD and 1KGP3. Finally, we pruned SNPs with linkage disequilibrium (*R*^2^ > 0.1) within 50 base pairs sliding window using PLINK.

The PCA was carried out on these processed data using Genome-wide Complex Trait Analysis (version 1.91.3beta)^51^ between 1) NARD and worldwide populations of 1KGP3, and 2) NARD and East Asians of 1KGP3, separately. We also applied the unsupervised ADMIXTURE algorithm (version 1.3) for the ancestry estimation. The optimal number of clusters was determined by comparing *K* values with cross-validation error rates (Supplementary Fig. 6). The number of ancestries was varied from *K* = 2 to *K* = 10. The results of ADMIXTURE analysis were visualized by Genesis (see URLs).

### Imputation

For the imputation panel, the singleton variants in NARD were excluded, because they are difficult to be imputed. To combine NARD and 1KGP3 panels, we used the same approach as UK10K and IMPUTE2 (See URLs); NARD-specific variants were imputed into 1KGP3 using Minimac3 (version 2.0.1) and vice versa, then merged into single reference panel. In addition, the combined panel was re-phased by SHAPEIT3 with ‘--early-stopping’ and ‘--cluster-size 4000’ parameters, and we kept variants that are not located in LCR.

For imputation accuracy evaluation, we separately processed 113 KOR individuals and 106 Northeast Asian individuals (79 CHN and 27 JPN) from previous studies^14,18,19,28,29^ using the same pipeline of NARD (Supplementary Table 3). These are an independent cohort from NARD, and we discarded 16 individuals in KOR cohort who have relatedness. Unrelated sample selection was achieved by kinship estimation using KING^52^. Then, we extracted SNPs on sites of Illumina Omni 2.5M array and monomorphic sites were excluded. As a result, 1,345,511 and 1,372,408 autosomal SNPs remained as pseudo-GWAS panels of KOR and CHN+JPN cohorts, respectively.

We performed imputation using Minimac3 with five different types of reference panel. Imputation using HRC panel was performed at Michigan Imputation Server. Before imputation, the haplotypes of individuals in two cohorts were estimated using Eagle (version 2.3.2). After imputation, we extracted 4,352,936 and 4,969,024 SNPs in two cohorts, respectively, which were imputed by all of five reference panels and none of true genotypes are not present as missing. The squared Pearson correlation coefficients (*R*^2^) were calculated between the imputed dosages and the true genotypes and those values were aggregated into 11 MAF bins to measure the imputation accuracy.

### IBD analysis

The shared IBD segments between two individuals were identified using RefinedIBD (version 12Jul18.a0b)^26^. To evaluate the effect of the re-phasing approach, we used the original haplotypes phased without reference panel and the re-phased haplotypes combined with 1KGP3 panel, separately. The short gaps and breaks (> 0.6 cM) between IBD segments were discarded using merge-ibd-segments utility program.

### URLs

Population statistics, http://www.worldometers.info/world-population

NARD imputation server, https://nard.macrogen.com/

Michigan Imputation Server, https://imputationserver.sph.umich.edu/

gnomAD, http://gnomad.broadinstitute.org/

DRAGEN, http://edicogenome.com/dragen-bioit-platform/

GATK’s best practice, https://software.broadinstitute.org/gatk/best-practices/

Picard Tools, https://broadinstitute.github.io/picard/

Genesis, http://www.bioinf.wits.ac.za/software/genesis/

NARD MAF data, https://nard.macrogen.com/download/NARD_Annovar.zip

Procedure for merge reference panels from IMPUTE2, https://mathgen.stats.ox.ac.uk/impute/merging_reference_panels.html

1KGP3 Illumina Omni 2.5M array data, ftp://ftp.1000genomes.ebi.ac.uk/vol1/ftp/release/20130502/supporting/hd_genotype_chip/ALL.chip.omni_broad_sanger_combined.20140818.snps.genotypes.vcf.gz

## Acknowledgements

We thank the members of GenomAsia100K Consortium for the discussion and assistance with the manuscript preparation. F.M. is supported by Japan Agency for Medical Research and Development (AMED) under grant numbers JP18kk0205008h0003, JP16ek0109070h0003, and JP18ek0109283h0001. K.-W.L. is funded by Research Grant Council, Hong Kong (Theme-based Research Scheme: T12-401/13-R).

## Author contributions

J.-S.S., C.K., S.S., J.-I.K., F.M., and B.C. designed the project. S.-K.Y., C.-U.K., H.L.K., and S.K. wrote the manuscript. S.-K.Y., C.-U.K., H.L.K., and S.K., performed the bioinformatic analysis. C.-U.K. and N.K. built imputation server. J.-Y.S. performed the library preparation and the next-generation sequencing. K.-W.L. and J.S.Y. contributed to the data interpretation.

## Competing interests

The authors declare no competing interests.

