## Supplementary Figure 1 for "Whole-genome reference panel of 1,781 Northeast Asians improves imputation accuracy of rare and low-frequency variants"

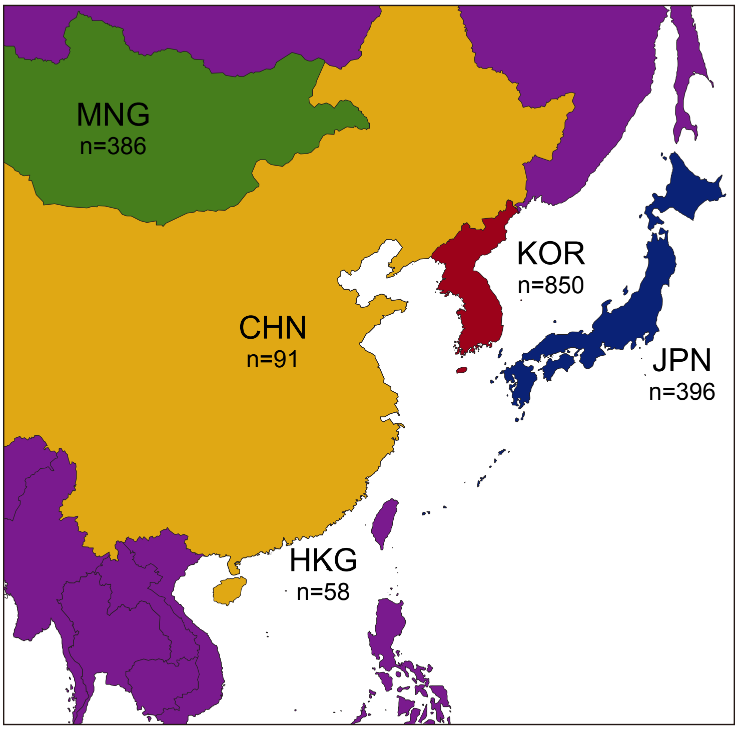


**Supplementary Figure 1. Geographic map of the study area in the Northeast Asian Reference Database (NARD).**
