## Supplementary Figure 2 for "Whole-genome reference panel of 1,781 Northeast Asians improves imputation accuracy of rare and low-frequency variants"

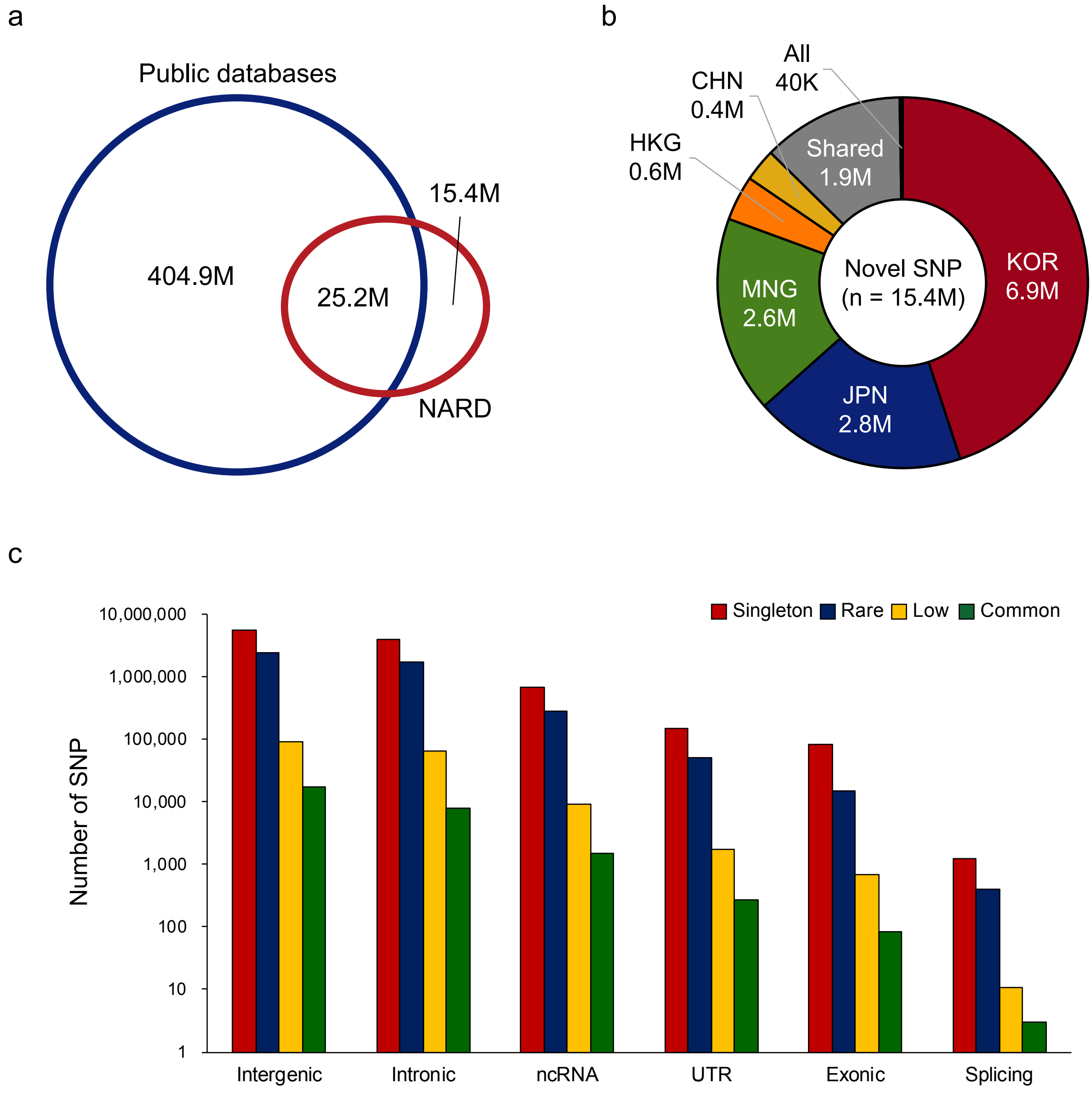


**Supplementary Figure 2. Novel variant statistics. a,** Number of novel SNPs that were not identified in elsewhere. Public databases include Kaviar, gnomAD (2.1.1 release), and dbSNP150. **b,** Distribution of novel SNP per population. Novel SNPs found in multiple and all populations were included in ‘Shared’ and ‘All’, respectively. c, Distribution of novel SNPs based on RefSeq gene definition.
