## Supplementary Figure 3 for "Whole-genome reference panel of 1,781 Northeast Asians improves imputation accuracy of rare and low-frequency variants"

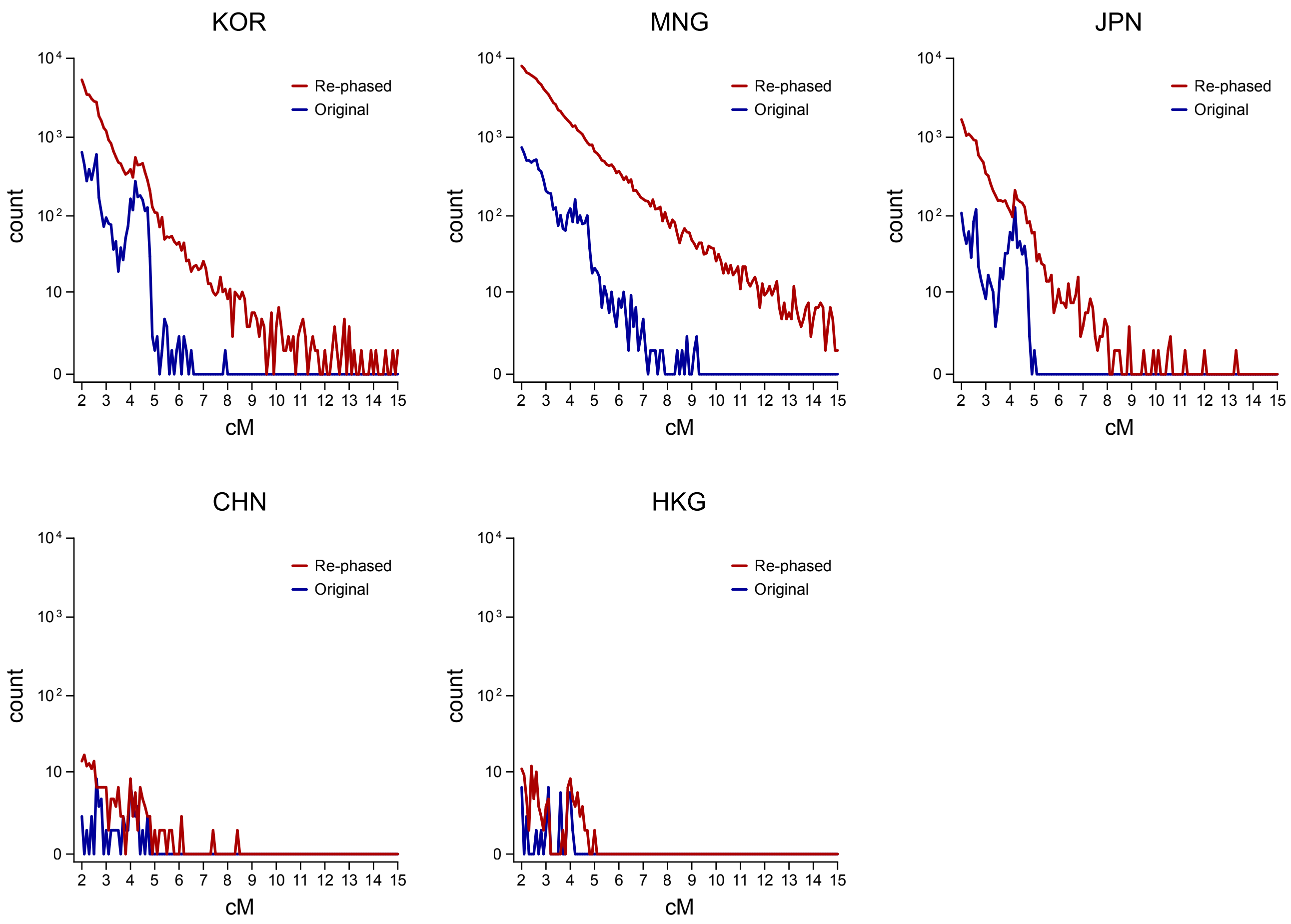


**Supplementary Figure 3. Length distribution of shared IBD tracts between the two individuals in each population.** Distributions of shared IBD tracts which were computed using the original and the re-phased haplotypes were displayed separately. Lengths of the shared IBDs are different by populations, but consistently increased in the re-phased haplotypes across all.
