## Supplementary Figure 4 for "Whole-genome reference panel of 1,781 Northeast Asians improves imputation accuracy of rare and low-frequency variants"

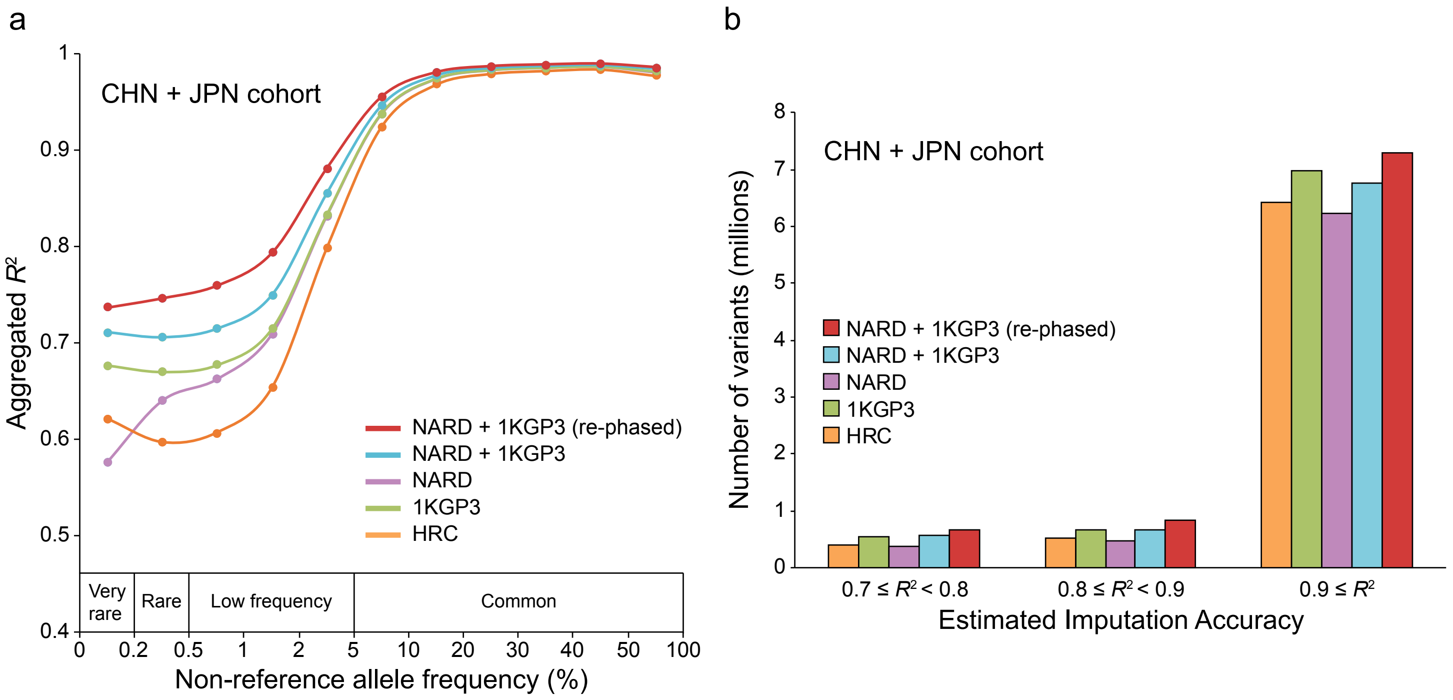


**Supplementary Figure 4. Imputation performance evaluation. a,** Imputation accuracy assessement using five different reference panels. The pseudo-GWAS panel of 106 Northeast Asian individuals was used for imputation. The *x*-axis represents MAF of 2,093 Northeast Asian individuals from NARD and 1KGP3. The *y*-axis represents the aggregated $R^2$ values of SNPs which were calculated by the true genotypes and the imputed dosages. Only SNPs that were imputed across all panels were used for the aggregation of $R^2$ values. **b,** Number of imputed SNPs as a function of the estimated imputation accuracy and the types of imputation panel. This result was generated based on the $R^2$ values that were estimated by Minimac3.
