## Supplementary Figure 5 for "Whole-genome reference panel of 1,781 Northeast Asians improves imputation accuracy of rare and low-frequency variants"

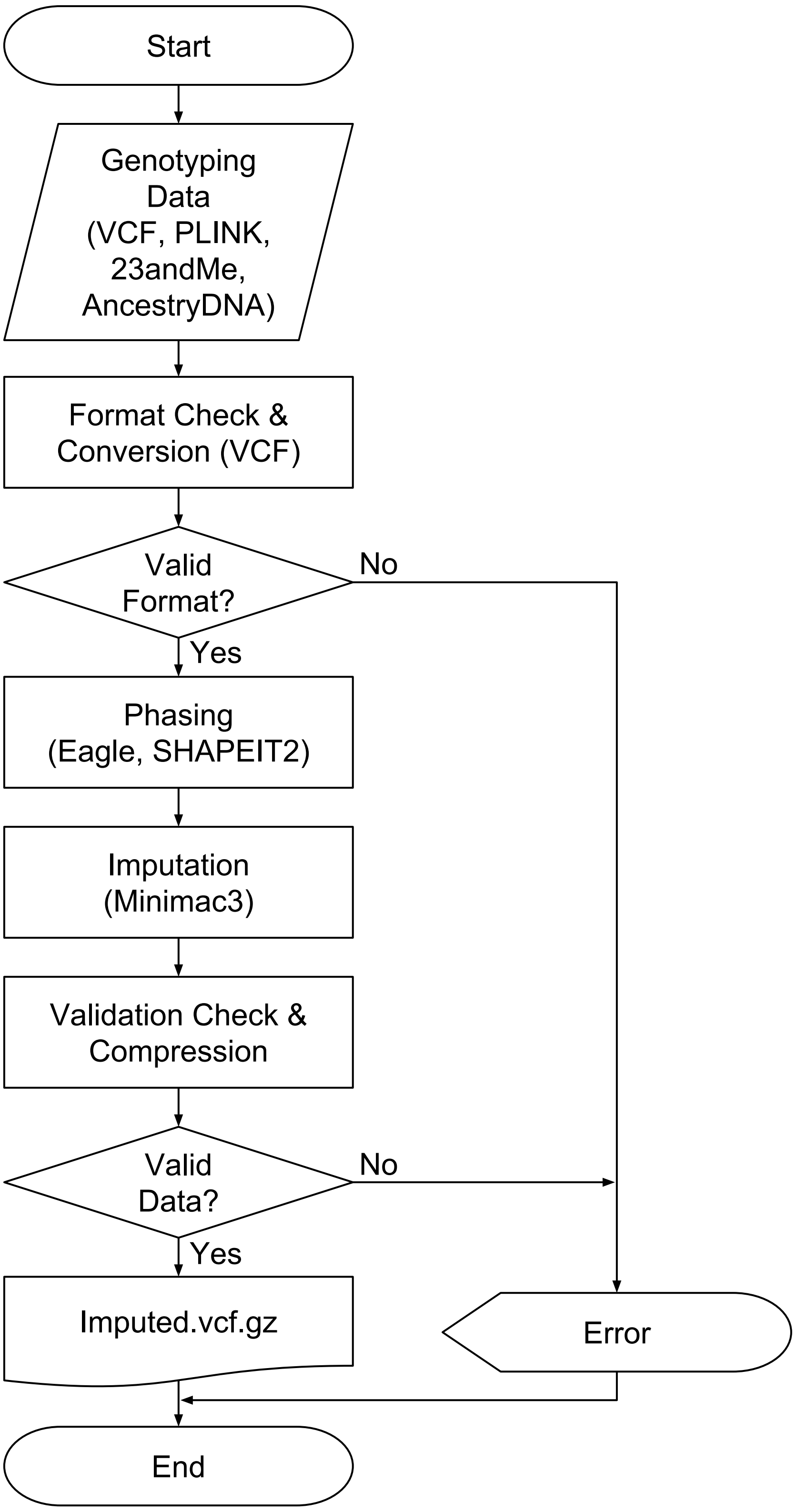


**Supplementary Figure 5. The flow chart of the pipeline consisting of four major steps for NARD imputation server.**
