## Supplementary figures and images for "Whole-genome reference panel of 1,781 Northeast Asians improves imputation accuracy of rare and low-frequency variants"

### Supplementary Figure 6

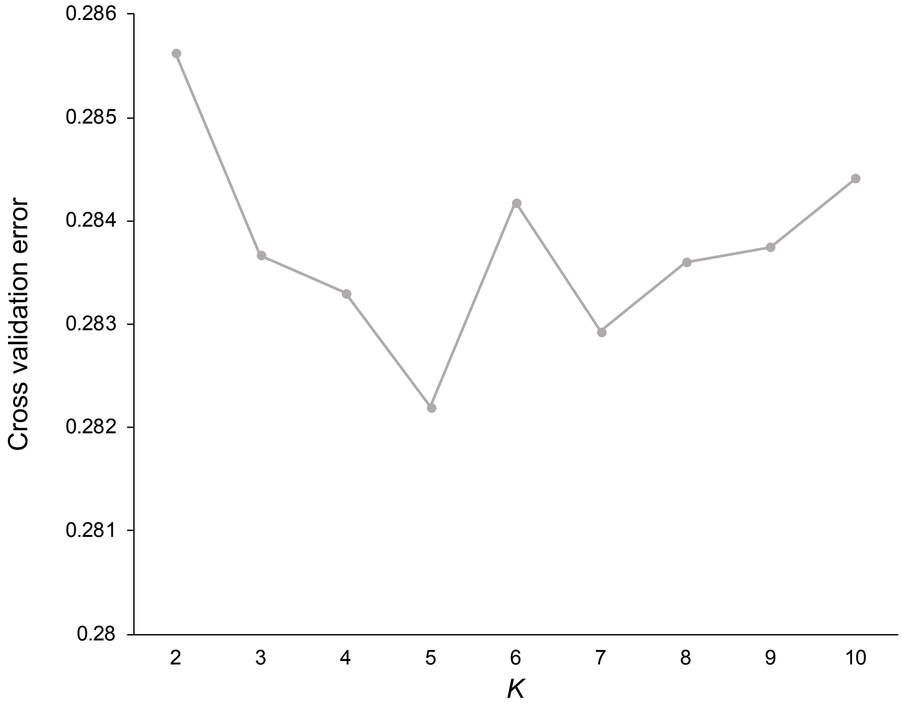


**Supplementary Figure 6. The cross-validation error inferred by ADMIXTURE algorithm.**
